# Adipocyte progenitors are primary contributors to the disrupted epithelial niche that is sustained following abrupt mammary gland involution

**DOI:** 10.1101/2024.09.08.611930

**Authors:** Prashant Nuthalapati, Dibyo Maiti, Sharon A. Kwende, Subhajit Maity, Dun Ning, Purna A. Joshi

**Affiliations:** Department of Biological Sciences, The University of Texas at Dallas, Richardson, TX 75080

## Abstract

A short duration of breastfeeding is a risk factor for the development of high-mortality, postpartum, triple-negative breast cancer. The intrinsic properties of cancer-initiating epithelial cells that persist following breastfeeding cessation and mammary gland remodeling are poorly understood. Previously, we showed that Platelet-Derived Growth Factor Receptor alpha (PDGFRα)-expressing stromal mammary adipocyte progenitors (MAPs) differentiate into epithelial luminal progenitors in the adult gland. In the current study, we demonstrate that MAP-derived luminal progenitors retain a mesenchymal transcriptomic signature. In an abrupt involution model that mimics a short breastfeeding duration, MAP-derived luminal progenitors persist and dominate luminal epithelia, undergoing transcriptomic alterations that signify a distinct ferrometabolic state linked to cancer. Concurrently, MAPs adopt an alternative interferon-mediated profibrotic and invasive stromal fate. Our work uncovers MAPs to be the primary cellular origin of a pathological stromal and epithelial microenvironment following abrupt involution, presenting a potential therapeutic target in postpartum breast cancer.

## Introduction

Epidemiological studies have demonstrated a complex interplay of factors such as parity, age, and breast-feeding patterns that influences pregnancy-associated breast cancer risk^1^. In particular, the 10-year postpartum period poses a window of heightened breast cancer risk and poor prognosis^2^. Further, a shorter duration of breastfeeding is associated with more aggressive, metastatic triple-negative, or basal-like breast cancer^3,4^. Probing fundamental alterations in the mammary gland following pregnancy and lactation is crucial for deconvoluting the cellular and molecular mechanisms behind pregnancy-associated breast cancer.

The mammary gland is an organ unique to mammalian species, with the primary function of providing nourishment for offspring via the secretion of milk. It comprises a parenchymal, bilayered epithelium that consists of outer basal myoepithelial cells lining an inner layer of luminal secretory cells^5,6^. The luminal population is significantly heterogenous^7,8^, and luminal progenitor cells that give rise to milk-producing luminal cells are putative targets in the pathogenesis of triple-negative breast cancers^9,10^. The epithelial ductal structure is embedded in a stromal fat pad that comprise adipocytes, fibroblasts, blood vessels, and tissue-resident immune cells. Throughout postnatal life, the mammary gland undergoes dynamic and significant alterations, functionally differentiating to form milk-producing lobuloalveoli during pregnancy and lactation. After weaning of offspring, the gland involutes—a physiologically normal process, akin to injury, that remodels the epithelial-dense, lactational gland back to a non-secretory state.

Involution is characterized by two distinct stages wherein the first reversible phase, when milk secretion can be reinitiated, transitions to an irreversible phase after 48 hours characterized by alveolar collapse, extensive extracellular matrix (ECM) remodeling, immune infiltration and adipocyte repopulation^11-14^. In murine models, a vast majority of the lactational mammary epithelium dies as part of this injury-like tissue remodeling process. However, unique properties of the surviving epithelial cells that evade cell death post-involution are not well known. Deciphering intrinsic characteristics of such cell populations is important for devising strategies to intercept the cellular origins of postpartum breast cancer. We previously identified PDGFRα-expressing mammary adipocyte progenitors (MAPs) that generate luminal progenitors in the mammary gland, especially contributing to the expansion of lobuloalveoli during hormone exposure and pregnancy^15^. Here, we explored the dynamics of MAP-derived luminal cells following pregnancy in an abrupt involution model^16^ which mimics a shorter duration of breastfeeding in humans. Using high resolution single-cell transcriptomic and phenotypic analyses, we show that MAP-descendants retain their mesenchymal features in the epithelium, comprise the majority of the luminal subset that remains post-involution, and adopt an aberrant stromal cell fate.

## Results

### Mesenchymal-like subset of luminal progenitors in the adult mouse mammary epithelium

To understand the contribution of MAPs and their progeny, we leveraged the high-fidelity *Pdgfrα* genetic reporter strain, *Pdgfrα*^*H2B-eGFP*^, which harbors a H2B-eGFP fusion fluorescent protein controlled by the endogenous *Pdgfrα* promoter^17^. The work of multiple labs across organ systems has shown the H2B-eGFP fusion protein to be stable, even in the absence of *Pdgfra* transcript, and thus is detectable in GFP^lo^ *Pdgfra*^-^ postmitotic cellular descendants in addition to the parent GFP^hi^ *Pdgfra*^+^ cells^15,18-21^.

Single-cell RNA sequencing (scRNA-seq) was performed on total GFP^+^ (both GFP^hi^ and GFP^lo^) cells sorted from mammary glands harvested from a nulliparous female *Pdgfrα*^*H2B-eGFP*^ mouse at a resting estrous phase (non-diestrus) (**Figures 1A and S1**). To elucidate the cellular identities of GFP-labelled cells in the context of the entire mammary microenvironment, data was integrated with a previously published scRNA-seq dataset^22^ of total cells from nulliparous glands of the same strain (**Figure 1A-F**). Following quality control and doublet removal, 33,715 unique barcodes were used for downstream analyses. Expected cell populations were observed including MAPs/fibroblasts, luminal epithelial cells, basal epithelial cells, endothelial cells, and immune cells (**Figures 1B and 1C**). Our GFP^+^ MAP-enriched population was well represented in the total fibroblast cluster in the Bach et al dataset (**Figure 1D**). GFP^+^ cells are also represented in endothelial, immune, basal and luminal progenitor subsets (**Figure 1D**).

**Figure 1.**
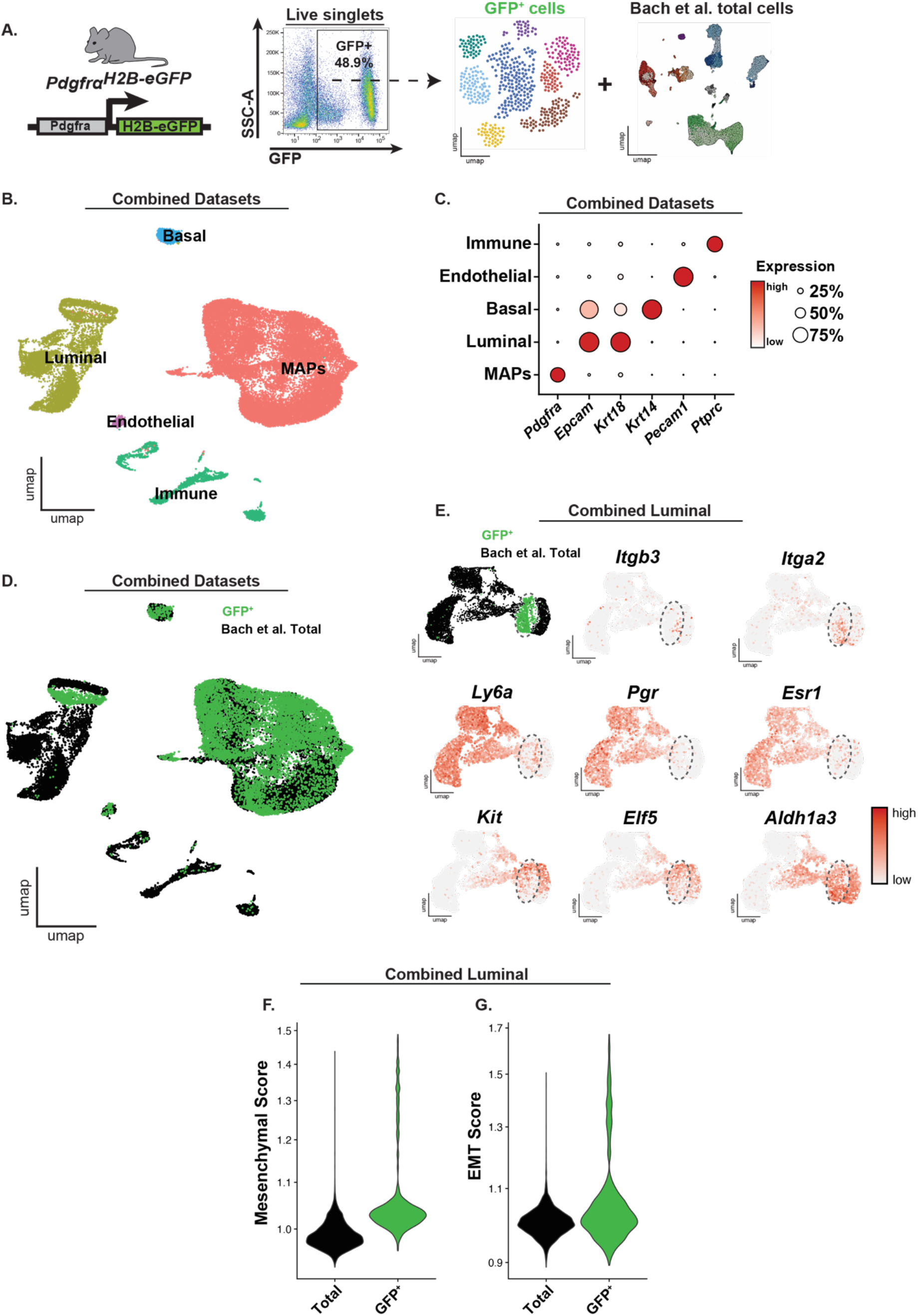
Mesenchymal-like subset of luminal progenitors in the adult mouse mammary epithelium. **(A)** Live GFP^+^ mammary cells from an adult nulliparous (non-diestrus) *Pdgfra*^*H2B-eGFP*^ female reporter mouse were sorted, sequenced, and merged *in silico* with Bach et al total, adult nulliparous mammary gland cells of the same background. **(B)** Uniform manifold approximation and projection (UMAP) of GFP^+^ mammary cells merged with Bach et al total mammary cells representing 33,715 transcriptomic profiles in MAP, luminal epithelial, basal epithelial, immune, and endothelial clusters. **(C)** Dot plot of canonical transcriptomic markers to identify broad cell populations. Size of circle indicates percent of cells with expression, and intensity of red indicates average expression. **(D)** UMAP of GFP^+^ mammary cells (green) merged with Bach et al total mammary cells (black) visualized by dataset. **(E)** Luminal cells represented in UMAPs, overlayed with curated luminal subpopulation markers. **(F)** Violin plot of mesenchymal gene signature comparing GFP^+^ vs total luminal cells. **(G)** Violin plot of epithelial-mesenchymal transition gene signature comparing GFP^+^ vs total luminal cells.

Interestingly, GFP^+^ luminal progenitors represent a subset that is enriched for largely hormone receptor-negative (devoid of *Pgr, Esr1, Ly6a*) alveolar progenitors predominantly expressing *Itga2* (CD49b), *Kit, Elf5* and *Aldh1a3* (**Figure 1E**).

Since previous work established the differentiation potential of PDGFRα^+^ MAPs to generate luminal progenitors by combined lineage tracing and transplantation assays^15^, we speculated whether MAP luminal progeny may retain features of its prior mesenchymal lineage. To test conserved mesenchymal fibroblast-like features, we curated a gene signature calculated from genes differentially expressed in fibroblasts compared to all other cell types in the Bach et al dataset (**Table S1**). We find that the GFP^+^ luminal cells have a higher mesenchymal gene score compared to total luminal cells from the published dataset (**Figure 1F**). Furthermore, GFP^+^ luminal cells possess a greater gene signature score ‘HALLMARKS_EPITHELIAL_MESENCHYMAL_TRANSITION’ obtained from the Mouse MSigDB Database ^23,24^ associated with epithelial-to-mesenchymal transition. Thus, a subset of luminal cells in the mammary gland derived from the MAP stromal lineage preserve transcriptomic features reminiscent of its mesenchymal origin.

### *Pdgfra* lineage descendants dominate the luminal compartment and generate a new stromal subset following abrupt involution

A shorter duration of breastfeeding can be recapitulated in abrupt involution murine models when weaning is artificially induced at the peak of lactation^25,26^. Adult *Pdgfrα*^*H2B-eGFP*^ female mice were analyzed at nulliparous (NP), mid-pregnancy (MP) at day 12.5 of gestation, and post-involution (PI) at day 14 following abrupt involution (**Figure 2A**). Whole mounts validated that nulliparous mice had fully developed ductal trees, mid-pregnant mice displayed lobuloalveologenesis, and post-involution mice had regressed, but remnant ductal and alveolar structures (**Figure 2B**). By immunofluorescence and confocal microscopy, we observed GFP^hi^ cells in the stromal compartment, while GFP^lo^ cells were detected in epithelia, primarily localizing to residual alveolar structures in post-involution glands (**Figures 2C and 2D)**.

**Figure 2.**
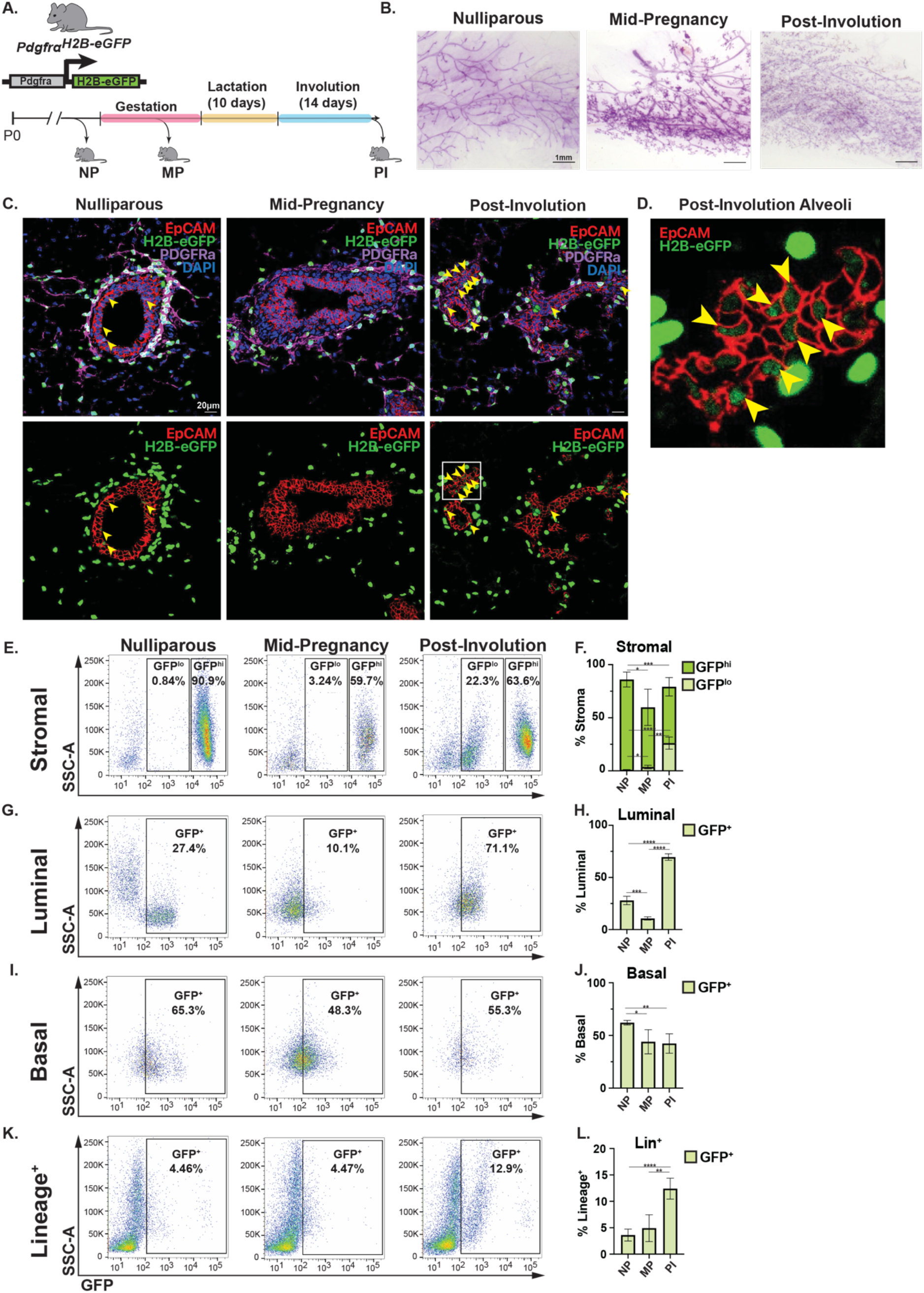
*Pdgfra* lineage descendants dominate the luminal compartment and generate a new stromal subset following abrupt involution. **(A)** Schematic of experimental protocol and timepoints: nulliparous (NP), mid-pregnancy d12.5 (MP), and post-involution d14 (PI). **(B)** Carmine alum-stained whole mount representative images, (n=3 per group; scale bar = 1mm). **(C)** Immunofluorescence images of *Pdgfrα*^*H2B-eGFP*^ mammary glands at the indicated experimental timepoints showing staining for EpCAM (red), PDGFRa (white), DAPI (blue), and unstained endogenous H2B-eGFP (green) (n=3 per group; scale bar = 20μm). White square denotes a representative post-involution remnant alveolar structure. **(D)** Digitally-zoomed image of field outlined in C, showing EpCAM (red) and H2B-eGFP (green) expression in a remnant alveolar structure of a post-involution mammary gland. **(E-L)** Representative flow cytometry and vertical bar plots of GFP-expressing cells in stromal, basal, luminal, and lineage^+^ (Ter119/CD31/CD45) mammary subpopulations of reporter mice (n=5 for NP, 3 for MP, and 4 for PI); bar plots depict the average percent of a subpopulation that expresses GFP (∗=*p*< 0.05, **=*p*<0.01, ***=*p*<0.001, ****=*p*<0.0001). Data is represented as mean ± SD.

By flow cytometry, we analyzed GFP^+^ cell proportions within distinct mammary subpopulations (**Figure S2**) using established cell surface markers ^8,27^ for segregating lineage^+^ (Lin^+^; CD45^+^CD31^+^Ter119^+^) and lineage^-^ (Lin^-^) luminal, basal and stromal subsets based on expression of CD49f and EpCAM. As expected, we detected a GFP^hi^ MAP-enriched population (85.1% ± 7%) primarily in the stromal compartment of nulliparous glands (**Figures 2E and 2F**). In mid-pregnancy, a decrease in the proportion of these GFP^hi^ cells is observed (56.3% ± 17.1%), and in post-involution, the GFP^hi^ population remains significantly lower (53.1% ± 8.7%). Interestingly, a GFP^lo^ stromal subset (26.1% ± 5.8%) emerged in post-involution (**Figures 2E and 2F**). Concordant with our previous study^15^, GFP^lo^ cells were seen in epithelial fractions, indicative of their origin from GFP^hi^ MAPs. In the luminal compartment, these GFP^+^ cell proportions changed from 27.9% ± 4.0% in nulliparous to 10.7% ± 1.6% in mid-pregnancy to a striking 69.6% ± 3.1% in post-involution, comprising the majority of the luminal population (**Figures 2G and 2H**). The basal compartment demonstrated a small, but significant decrease in the proportion of GFP^+^ cells from nulliparous (62.1% ± 2.2%) to mid-pregnancy (44% ± 11.5%) which was maintained following abrupt involution (42.4% ± 9.2%) (**Figures 2I and 2J**). The Lin^+^ compartment also exhibits a small and insignificant expansion in the proportion of GFP^+^ cells between nulliparous (3.6% ± 1.1%) and mid-pregnancy (4.9% ± 2.5%), but is significantly increased post-involution (12.4% ± 2.0%) (**Figures 2K and 2L**).

Overall, descendants of the *Pdgfra*^*+*^ cell lineage show dynamic shifts in the mammary microenvironment, characterized by the appearance of a new stromal subpopulation and the dominance of MAP-derived luminal cells following abrupt involution.

### Heterogeneity and cell fate decisions of MAPs are altered in the post-involution gland

We then sought to define alterations in the cellular lineage marked by the *Pdgfra* reporter following abrupt involution. As done for nulliparous mice, we conducted scRNA-seq on sorted live GFP^+^ cells from post-involution *Pdgfra*^*H2B-eGFP*^ mammary glands (**Figure S1**). After merging our GFP^+^ *Pdgfra* lineage post-involution dataset with our GFP^+^ nulliparous dataset, we observed MAPs, luminal progenitors, basal epithelial, endothelial and immune cell populations as detected in the nulliparous gland (**Figures 3A and 3B**). Subsequently, we isolated and subclustered the MAPs *in silico* to resolve MAP heterogeneity and differentiation dynamics within the stromal compartment (**Figure 3C**). In doing so, 8 clusters of MAPs emerged, each with distinct transcriptomic profiles (**Figure 3D**). When subjected to the unbiased, CytoTRACE differentiation hierarchy analysis that predicts developmental potential based on gene count^28^, the *Pi16*^+^ MAP4 cluster was the most developmentally immature subtype concordant with previous pan-tissue studies^29^ (**Figure 3E**). Interestingly, the differentiation hierarchy of MAPs was perturbed in post-involution, wherein the most differentiated *Fam13a*^+^ MAP3 subtype in the nulliparous gland became relatively less differentiated, and multiple MAP subtypes (*Rgcc*^+^ MAP1, *Ifit1*^+^ MAP6, *Csmd1*^+^ MAP8) increased in their differentiation potential, inferring a preference for alternative MAP cell states post-involution. Pseudotemporal ordering^30^ indicated a differentiation trajectory from the *Pi16*^+^ MAP4 progenitor subtype to the expanded MAP6 cluster in the post-involution gland (**Figure 3F**).

**Figure 3.**
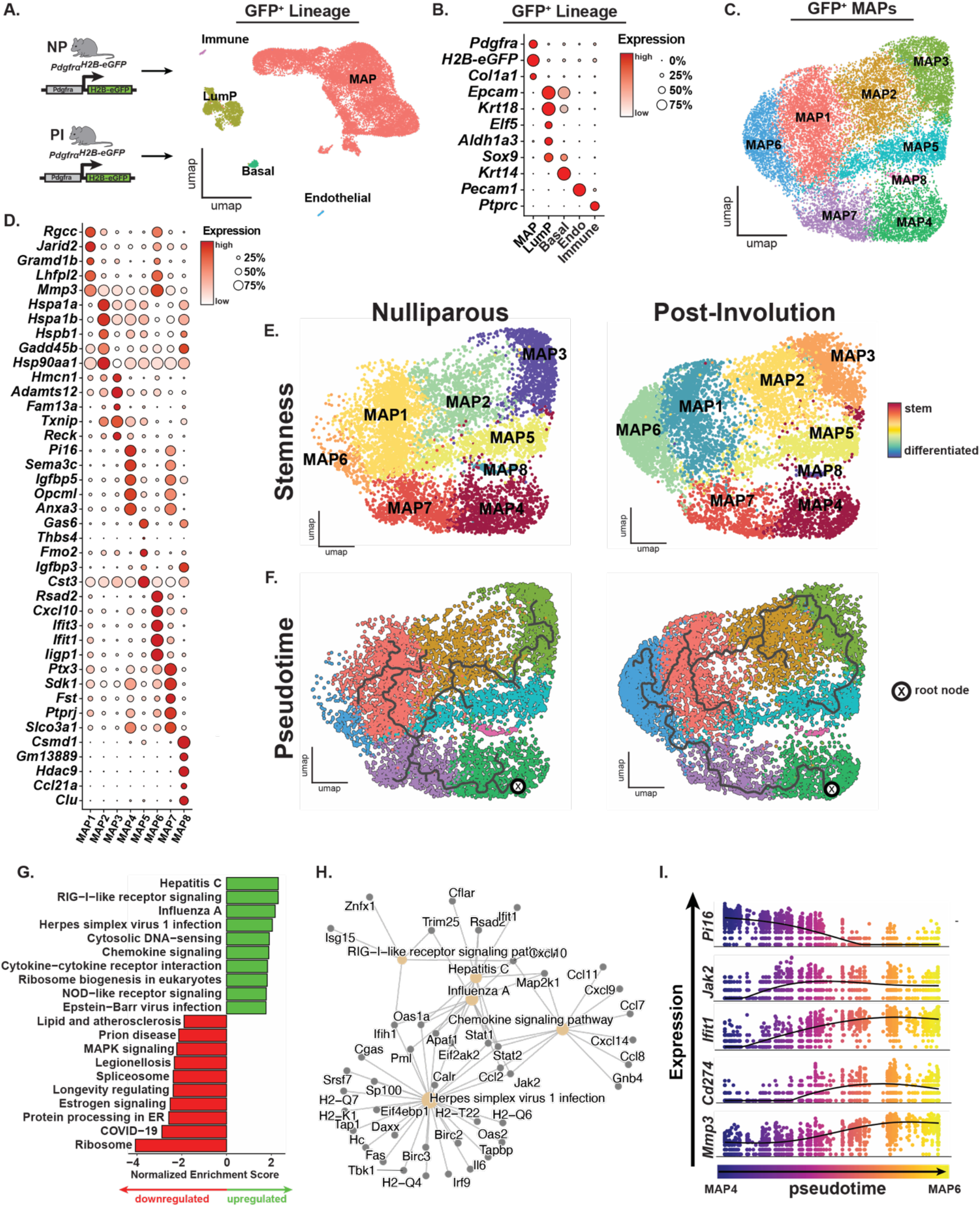
Heterogeneity and cell fate decisions of MAPs are altered in the post-involution gland. **(A)** Schematic depicting processing of GFP^+^ cells from nulliparous and post-involution *Pdgfrα*^*H2B-eGFP*^ reporter mammary glands for computational analysis. UMAP of merged dataset consisting of 19,215 transcriptomic profiles, represented in MAP, immune, luminal epithelial progenitor, basal epithelial, and endothelial clusters. **(B)** Dot plot of curated gene set to identify cell identities. Size of circle indicates percent of cells with expression, and intensity of red indicates average expression. **(C)** UMAP of GFP^+^ MAPs combined from nulliparous and post-involution. **(D)** Dot plot of top 5 differentially expressed genes in each cluster. Size of circle indicates percent of cells with expression, and intensity of red indicates average expression. **(E)** CytoTRACE score overlaid on UMAP of MAPs, split by nulliparous and post-involution. Red represents most stem, and blue represents most differentiated. **(F)** Pseudotime trajectory analysis of MAPs, split by nulliparous and post-involution. Root node, denoted by “X” was selected using most stem cluster by CytoTRACE. **(G)** Horizontal bar plot depicting top 10 upregulated and downregulated KEGG pathways in MAP6 compared to other MAPs, ordered by normalized enrichment score. **(H)** Network plot depicting gene-pathway relationships in the top 5 upregulated KEGG pathways, by *p*-value. **(I)** Gene expression of curated transcriptional targets along pseudotime of MAP4 to MAP6 in the post-involution dataset.

KEGG pathway analysis of the MAP6 cluster versus all other MAP clusters reveals significant upregulation of viral infection-associated pathways which comprise 5 of the top 10 upregulated pathways.

Additionally, a down regulation of lipid and atherosclerosis pathway in MAP6 suggests it to be a non-adipogenic cluster that assumes a cell fate opposing adipogenesis (**Figure 3G**). To better understand transcriptional drivers of MAP6 identity, top 5 most significant, and upregulated KEGG pathways in MAP6 were visualized with a network plot (**Figure 3H**). Consensus driver genes within these pathways included *Stat1, Stat2*, and *Jak2* (**Figure 3H**). Furthermore, the expression of interferon-induced protein with tetratricopeptide repeats 1 (*Ifit1*) implicates the IFN/JAK2/STAT1 axis^31^ in the induction of MAP6. The IFN pathway has been reported to stimulate PD-L1 expression in tumor-associated immunoregulatory fibroblasts^32^. Indeed, we observed an induction in the expression of *Cd274* (PD-L1), *Ifit1, Jak2* and termination in *Pi16* expression along the pseudotime trajectory from MAP4 to MAP6 (**Figure 3I**). The expression of *Mmp3*, a proteinase involved in mammary cancer^33,34^, was also enhanced during this transition suggestive of the invasive state of MAP6 (**Figure 3I**). Together, our analyses pinpoint critical shifts in MAP heterogeneity following abrupt involution where MAPs exhibit changes in cell state and differentiation potential, generating a MAP subpopulation reminiscent of tumor-associated fibroblasts.

### The *Pdgfra* lineage-derived luminal population undergoes ferrometabolic shifts post-involution

Next, we investigated single-cell transcriptomic changes in MAP-derived luminal progenitors given our observation of a sustained and dominant GFP^+^ subset in the luminal epithelium following abrupt involution in *Pdgfra* reporter mice. We sorted the luminal progenitor (LP) cluster *in silico* from our GFP^+^ nulliparous and post-involution scRNA-seq data in Figure 3A and upon subclustering, we uncovered dramatically altered heterogeneity (**Figure 4A**). Three luminal progenitor subsets were found with LP1 predominantly existing in the nulliparous gland. In abrupt involuted glands, LP2 and LP3 proportionally expand to over 75% of the GFP^+^ luminal compartment (**Figure 4B**). To analyze the transcriptomic signature of the various luminal progenitor subsets, we ascertained the top 5 genes that are specific to each cell population through differential gene expression analysis (**Figure 4C**). Interestingly, LP2 and LP3 specifically expressed lactation genes *Csn1s1* and *Spp1* compared to LP1. Previously, in natural involution models, the expression of the casein locus post-involution was attributed to changes in chromatin accessibility of the luminal progenitor population following lactation^7^. To determine whether MAP-derived GFP^+^ luminal cells also maintain this trend, we curated a lactational signature (*Csn1s1, Csn2, Csn1s2a, Csn1s2b, Csn3, Wap, Mfge8*) and found that MAP-derived luminal progenitors in post-involution possessed a higher lactation gene signature compared to those in nulliparous glands (**Figure 4D**). The presence of this lactation profile in post-involution luminal progenitors indicates their prior existence in the lactogenic epithelium, and their subsequent selection for survival in the apoptotic involuting gland.

**Figure 4.**
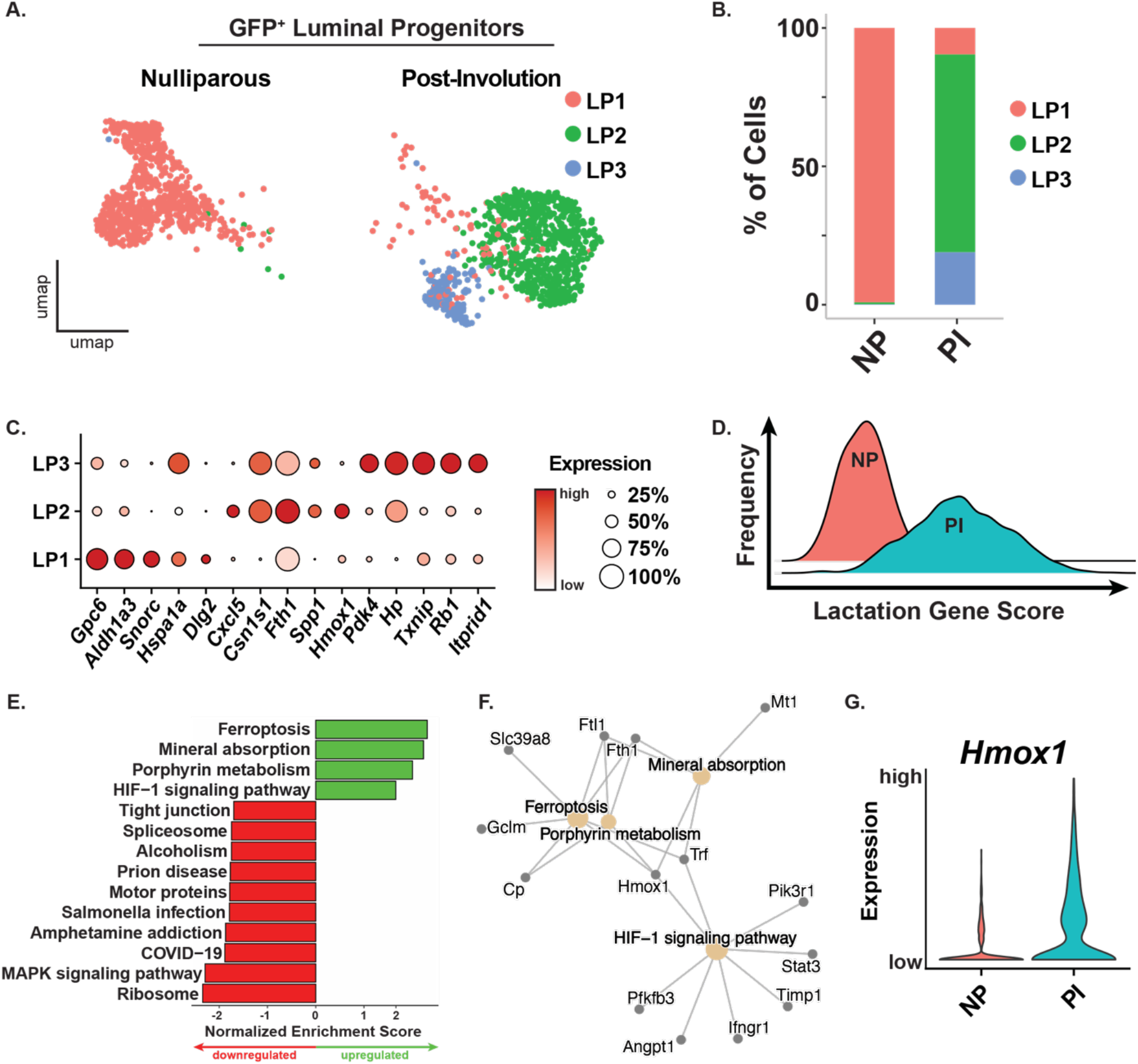
The *Pdgfra* lineage-derived luminal population undergoes ferrometabolic shifts post-involution. **(A)** UMAP of GFP^+^ luminal progenitor cells, split by nulliparous and post-involution states, represented in 3 luminal progenitor clusters. **(B)** Bar plot depicting cell proportion shifts following abrupt involution. **(C)** Dot plot representing top 5 differentially expressed genes per cluster. Size of circle indicates percent of cells with expression, and intensity of red indicates average expression. **(D)** Ridge plot depicting the expression of a curated lactation gene signature in post-involution compared to nulliparous GFP^+^ luminal cells. **(E)** Horizontal bar plot depicting upregulated (green) and downregulated (red) KEGG pathways in post-involution compared to nulliparous GFP^+^ luminal progenitors, ordered by normalized enrichment score. **(F)** Network plot depicting gene-pathway relationships in upregulated KEGG pathways. **(G)** Violin plot of gene expression levels of *Hmox1* in nulliparous and post-involution GFP^+^ luminal cells.

To elucidate the intrinsic properties of MAP-derived luminal progenitors, and the changes they undergo in adapting to involution-associated selection pressures, we analyzed the top upregulated and down regulated KEGG pathways between post-involution and nulliparous states. Notably, we find 4 key upregulated KEGG pathways, with 3 of 4 pertaining to mineral-associated pathways (**Figure 4E**). To delineate underlying gene-pathway relationships between upregulated pathways, we produced a network plot which revealed heme-oxygenase-1 (*Hmox1*), to be a common transcriptional driver that was linked to all pathways (**Figure 4F**). A primary function of HMOX1 is the catabolic degradation of heme into biliverdin, carbon monoxide and ferrous iron^35^. HMOX1 is known to be induced in response to oxidative stress and be protective against apoptosis^36-38^. Indeed, *Hmox1* is upregulated in post-involution MAP-derived luminal cells compared to their nulliparous counterparts (**Figure 4G**). Thus, luminal progeny of MAPs are preferentially sustained following lactation and involution with heightened expression of the ferrometabolic regulator *Hmox1* which may promote its survival under the extensive apoptotic conditions of involution.

## Discussion

The pathogenesis of breast cancer is inextricably linked to the underlying biology of the mammary gland, which undergoes immense changes throughout the female life course. The involution process triggered by milk stasis is known to generate a mammary niche that promotes breast cancer invasion^16^. Prior studies have placed significant emphasis on the intrinsic properties of epithelial cell populations, relegating stromal cells to only indirectly orchestrate normal and pathological processes in the mammary gland via paracrine mechanisms. During the gestational and lactational period, adipocytes in the stroma dedifferentiate into PDGFRα^+^ adipocyte progenitors as epithelial cells proliferate and dominate the gland to undergo functional differentiation. Following abrupt involution, both the stromal and epithelial niches are altered in concert, characterized by concurrent fibroblast activation, wound healing-like extracellular matrix deposition, immunoregulatory signaling, ductal hyperplasia, and altered mammary epithelial subpopulations^25,26^. In the stroma, a PDGFRα^+^ stromal population was previously shown to be an important driver of tumorigenesis and immunosuppression during involution^26^. In a prior study, we have demonstrated remarkable plasticity in the *Pdgfra*-expressing MAP lineage in the mammary gland highlighted by their capacity to generate epithelial progeny, predominantly luminal progenitors during pregnancy and sex hormone exposure^15^. The intrinsic properties and dynamics of this MAP-derived luminal population post-pregnancy and involution of the mammary gland was hitherto unknown.

In this study, we report that MAP-derived luminal cells in the mammary gland transcriptomically resemble an alveolar luminal progenitor that retains mesenchymal characteristics. We find that these cells persist beyond pregnancy and lactation, becoming the primary constituent of the luminal epithelium after abrupt involution. Significant changes in the stroma post-involution include the presence of a new MAP-derived stromal population, and single-cell studies unveil a new PD-L1 (*Cd274*) MAP cell state harboring fibrotic potential following abrupt involution in the mammary gland.

Postpartum breast cancer associated with a shorter duration of breastfeeding is particularly linked to the incidence of aggressive, metastatic triple-negative breast cancer^3,4^. These basal-like breast cancers are increasingly recognized to have roots in the luminal progenitor lineage^9,10^. A previous report indicated an increase in luminal progenitors following abrupt involution^25^. We now show that the MAP-derived luminal population with mesenchymal traits is primarily the population that persists following abrupt involution. This suggests that the chromatin accessibility of the previous mesenchymal MAP cell state persists even after mesenchymal to epithelial transition, potentially contributing to the plasticity associated with oncogenic and metastatic conditions. The precise nature of luminal epithelial cells that survive vast apoptotic remodeling in the involuting gland is poorly defined. Our work demonstrates that mesenchymal-like alveolar progenitors that survives involution have ferrometabolic properties with marked expression of *Hmox1* that confer cellular resistance to apoptosis and oxidative stress. These apoptotic-resistant mesenchymal-like alveolar progenitors are likely dominant cellular targets in post-partum breast cancer given their potential capacity to evade host anti-tumor apoptosis mechanisms. Findings from our study highlight the intrinsic plasticity of mammary adipocyte progenitors and their progeny which endow them with a unique capacity to adapt to injury-like stress conditions in an epithelial tissue niche.

## Methods

### Animal handling

*Pdgfrα*_*H2B-eGFP*_ mice were bred on the C57BL/6J background and were maintained with a 12-hour day, 12-hour night cycle on standard rodent diet. Female mice were randomly assorted to experimental groups. The nulliparous, virgin group was sacrificed between 10-15 weeks of age. For the mid-pregnancy group, a wild-type C57BL/6J male was placed in the cage overnight, separated the following morning, and the female was subsequently weighed every day to ensure weight gain due to pregnancy. On the 12^th^ day of pregnancy (mid-pregnancy), the female was sacrificed for mammary gland isolation. For the post-involution group, the male was kept with the female until visibly pregnant, then separated to ensure a single pregnancy. On the 10^th^ day following parturition, the pups were removed from the cage to initiate abrupt involution, and 14 days after weaning, the female was sacrificed for mammary gland analysis. Nulliparous and post-involution females were validated to be in a non-diestrus stage of the estrus cycle via vaginal cytology. All animal handling was done in accordance with protocols approved by the Institutional Animal Care and Use Committee (IACUC) at the University of Texas at Dallas. To ensure consistency in downstream analysis, thoracic mammary glands were used for flow cytometry. Inguinal glands were used for mammary tissue whole mounts and immunofluorescent staining.

### Mammary tissue whole mounts

Immediately upon dissection, inguinal glands were spread on a positively charged glass slide and placed in Carnoy’s fixative for 24 hours at room temperature. Following fixation, glands were rehydrated in decreasing concentrations of ethanol (70 to 0%) in distilled water and were stained overnight with carmine alum. Stained glands were dehydrated in increasing concentrations of ethanol (70 to 100%), and cleared in xylene. Following clearing, glands were mounted in Permount and left to dry overnight. Whole mounts were imaged on the Olympus SZ61 stereo microscope.

### Immunofluorescent staining

Following dissection, inguinal glands were placed in 4% paraformaldehyde for 2 hours at room temperature for fixation. They were washed in PBS three times for 5 minutes each, placed in 30% sucrose solution overnight at 4°C, embedded in OCT, and stored in -80°C. Samples were cryosectioned at 10 μm thickness and slides were stored in -80°C.

For immunofluorescence staining, sections were blocked in 5% normal donkey serum/1% BSA/0.2% Triton in PBS at room temperature for one hour. Primary antibodies were diluted in blocking solution without Triton and left to stain overnight at 4°C. Next day, sections were washed three times for five minutes in PBS and were subsequently incubated with secondary antibodies for 1 hour at room temperature. Sections were then rinsed in PBS, and mounted in ProLong Gold Anti-fade reagent with DAPI nuclear stain. Stained slides were stored at 4°C.

Primary antibodies used were goat anti-PDGFRα (R&D systems, Cat.# AF1062) and rabbit anti-EpCAM (Abcam, Cat.# ab71916). Secondary antibodies used were anti-goat and anti-rabbit antibodies conjugated to AlexaFluor 647, AlexaFluor Cy3. Reporter H2B-eGFP signal was detected with endogenous fluorescence. Experimental groups were stained in bulk at the same time with the same reagents to ensure comparability. Immunostained sections were imaged on an Olympus FV3000RS Confocal Laser Scanning Microscope at 40X. Five fields were imaged per tissue section and experimental groups were imaged in triplicate.

### Tissue dissociation, flow cytometry and FACS

Digestion solution was prepared from a 3:1 collagenase:hyaluronidase solution in DMEM:F12 and warmed at 37°C. Immediately after dissection, both pairs of thoracic glands were briefly rinsed in PBS on ice and placed in warmed digestion solution for 2.5 hours at 37°C. Following digestion, glands were vortexed to fully dissociate tissues. The digestion was stopped with Hank’s balanced salt solution with 2% FBS. Tissues were centrifuged, and red blood cells were lysed with ammonium chloride solution. Following red blood cell lysis, tissues were incubated in 0.25% trypsin for 2 min, 5 mg/ml dispase with 0.1 mg/ml DNase for 2 min, and filtered through a 40μm mesh to generate a single cell solution.

Following dissociation, cells were stained with PE-Cy7 antibodies conjugated to CD45 (Ebioscience; Cat.# 25-0451), CD31 (Ebioscience, Cat.# 25-0311) and Ter119 (Ebioscience, Cat.# 25-5921) for immune, endothelial, and erythrocytes respectively. Jointly, this is referred to as lineage^+^ (Lin^+^) cells. Anti-CD49f/APC (R&D, Cat.# FAB13501A) and anti-EpCAM/APC-Cy7 (Biolegend, Cat.# 118217) was used to identify Lin^-^ stromal and epithelial subpopulations. Dead cells were excluded with Zombie UV viability dye (Biolegend, Cat.# 423107). GFP endogenous fluorescence was detected without staining. Flow cytometry was performed using a Fortessa cell analyzer and sorting using a FACSAria Fusion (BD) with FACSDiva software (BD), and FlowJo software was used for downstream analysis (Tree Star, Inc).

### Library preparation, sequencing, and preprocessing

Live GFP^+^ cells were FACS-sorted using the FACSAria Fusion (BD) from nulliparous and post-involution *Pdgfrα*^*H2B-eGFP*^ mice. A Fluorescence Minus One (FMO) control using non-reporter wild-type cells was used to determine the GFP^+^ gate for sorting. Sample preparation was conducted individually and in accordance with the 10X Chromium single-cell RNA sequencing kit. Sequencing for both samples was done together, minimizing batch effects, with the Illumina Nextseq 2000 instrument with the P2 100bp flow cell. Sequencing read was performed as follows: Read1: 28 cycles, Read2: 90 cycles, Index1: 10 cycles, Index2: 10 cycles. Sample demultiplexing, alignment, and filtering was performed with the Cell Ranger Cloud Analysis Software (v5.0.1) to a custom mm10 reference genome with the *H2B-eGFP* transgene annotated under “*GFP_new_id*”. In the nulliparous sample, 9,322 unique barcodes were recovered, and 10,939 unique barcodes were recovered in the post-involution sample.

Following library preparation, sequencing, and alignment, data was processed using Seurat (v4.4.0)^39^. Genes *Gm42418* and *AY036118* were removed due to potential rRNA contamination because of sequence overlap with the rRNA element Rn45s. Barcodes with greater than 8% mitochondrial reads were filtered to exclude low-quality cells. Doublets were computationally filtered with DoubletFinder (v2.0.4)^40^ in accordance with 10X multiplet rates. In total, 19,215 unique barcodes (9,121 in the nulliparous sample, and 10,094 barcodes in the post-involution sample) were ascertained to represent viable single cells and utilized in downstream analysis.

### Single-cell RNA sequencing analysis

Analysis was conducting using standard Seurat protocols in R Studio. To ascertain cell profiles of MAPs and *Pdgfra*-lineage descendants in the nulliparous mouse (**Figure 1A**), data was merged prior to normalization with pre-processed Bach et al. data^7^ using the harmony package (v1.2.0)^41^ due to observed batch effects.

To compare cell profiles of MAPs and *Pdgfra*-lineage descendants in nulliparous versus post-involution, data was merged prior to normalization without harmony (**Figure 3A**). Unsupervised hierarchical differentiation states and stemness were predicted with CytoTRACE (v0.3.3)^28^. Pseudotime trajectory inference was performed with the monocle3 package (v1.3.4)^30^. The root node was selected using the most stem cluster according to CytoTRACE. Differentially expressed genes (adjusted-*p*<0.05) were found using the Seurat FindMarkers function. KEGG pathway enrichment analysis was performed and visualized using the clusterProfiler package (v4.10.1)^42^.

### Statistical analyses

Independent mice, denoted by *n*, were pooled from independent experiments. Flow cytometry data was analyzed with FlowJo (v10.8.1) and visualized using GraphPad Prism (v10.0.0), reported as mean ± standard deviation (SD). Comparison of data between two groups was made using Student’s *t*-test (two-tailed). Statistical significance is recognized at *p*<0.05.

## Data availability

Processed and raw UMI count matrices will be deposited in the NCBI Gene Expression Omnibus. Code will be made available on GitHub. Pre-processed Bach et al data was downloaded directly from the authors’ (http://marionilab.cruk.cam.ac.uk/BRCA1Tumourigenesis) site to maintain integrity with previously published analyses.

## Acknowledgements

This work was supported by UT Dallas Startup funds to P.A.J. We acknowledge the UT Dallas Laboratory Animal Resource Center for assistance with animal care, the UT Dallas Flow Cytometry Core for cell sorting, the UT Dallas Genome Core for single-cell RNA sequencing, and the UT Dallas Cyberinfrastructure & Research Services Department for high-performance computing access.

## Author contributions

P.N. conceived the study, performed computational analyses, wet-lab experiments, and wrote the manuscript. D.M., S.A.K., and S.M. assisted with wet-lab experiments. D.N. assisted with colony management and animal handling. P.A.J. conceived, directed the study, and wrote the manuscript with P.N.

