## Supplemental Data for "Adipocyte progenitors are primary contributors to the disrupted epithelial niche that is sustained following abrupt mammary gland involution"

**Nuthalapati et al.**

**Supplemental Information**

### Supplemental Figure 1

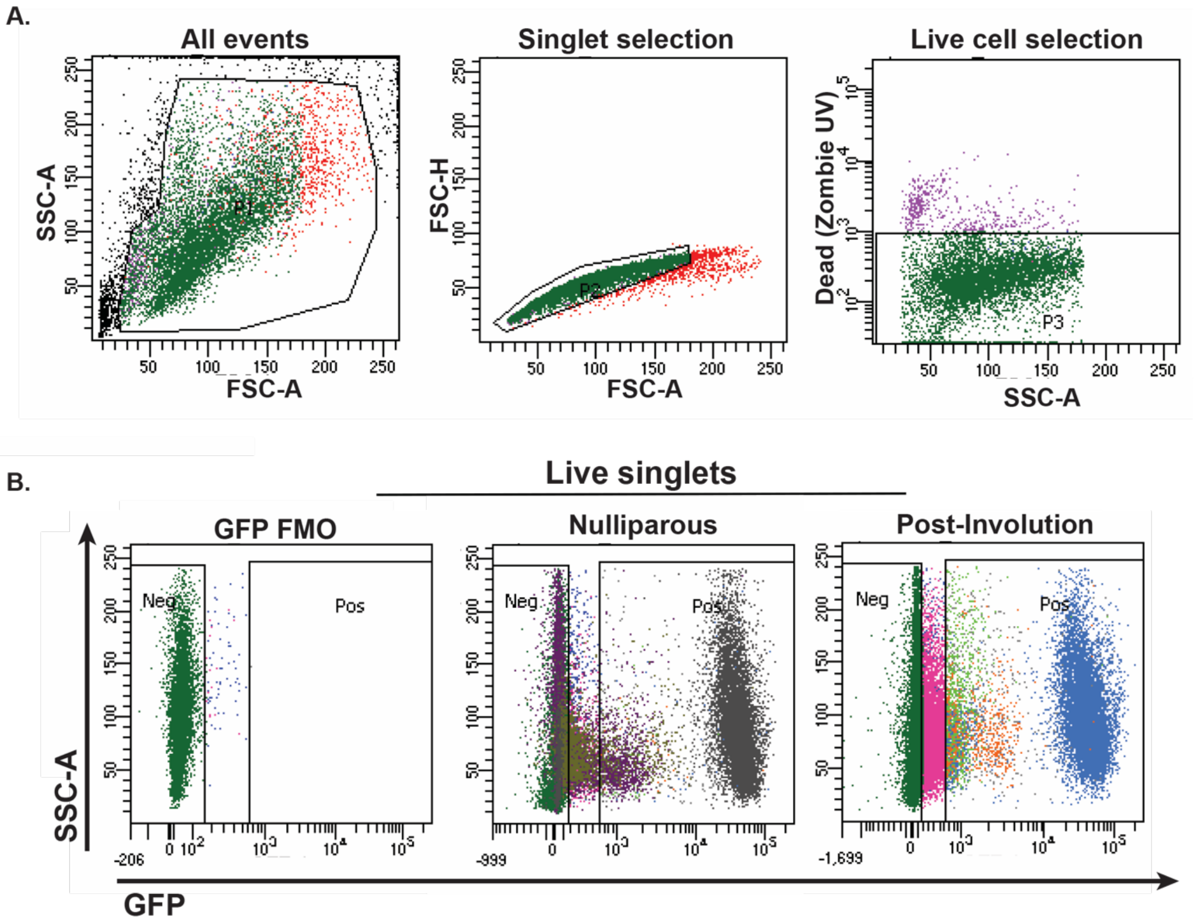


**Figure S1. Sorting strategy for single-cell RNA sequencing samples. (A)** Sorting strategy used to isolate live singlets for single-cell RNA sequencing. **(B)** Sort plots for single-cell RNA sequencing samples. Non-reporter wild-type Fluorescence Minus One (FMO) control was used to determine the GFP$^{+}$ gate for sorting.

### Supplemental Figure 2

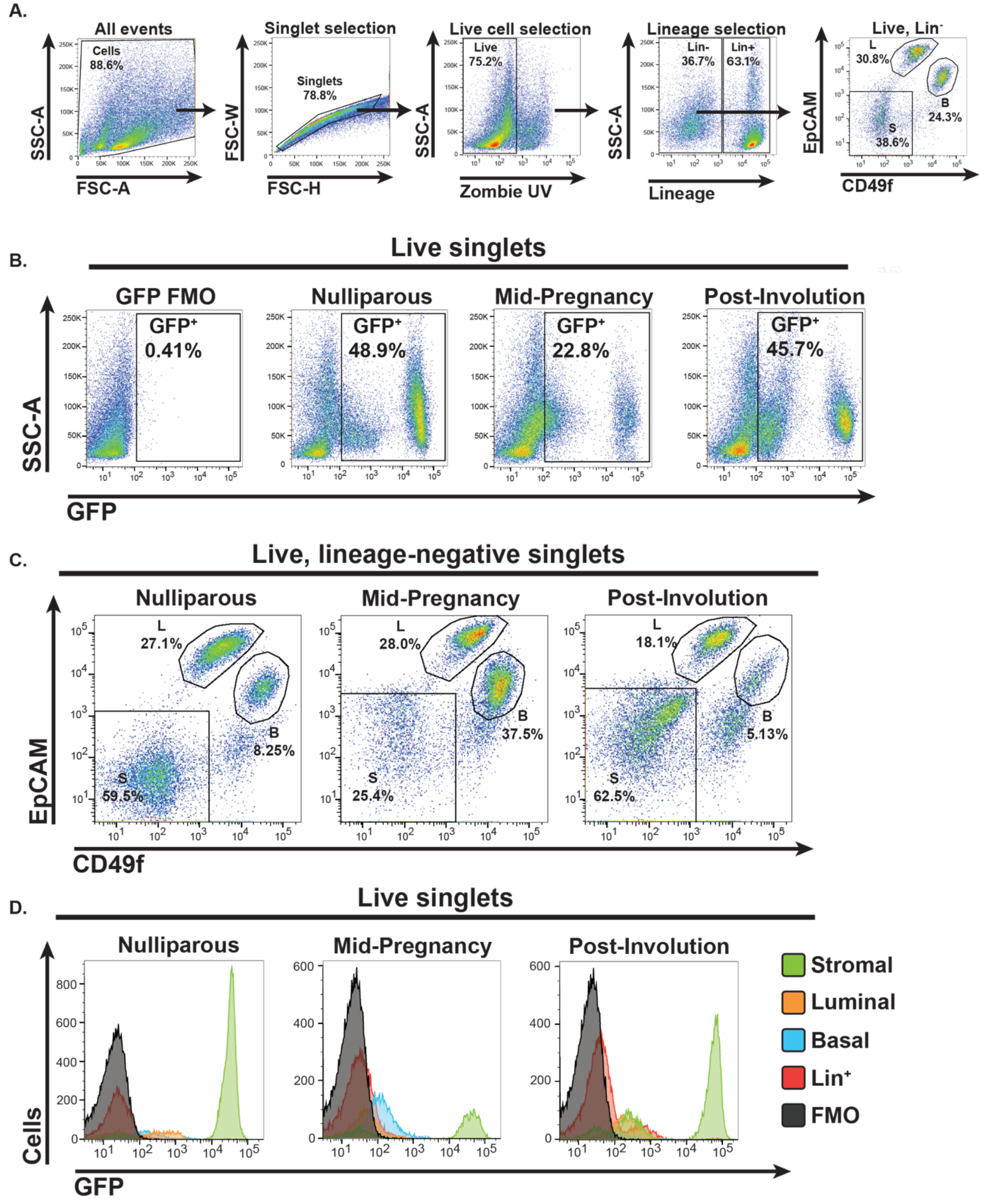


**Figure S2. Gating strategy for flow cytometry analysis. (A)** Gating strategy used to identify single, live, and lineage (CD45/CD31/Ter119)-negative cells. **(B)** GFP expression in total, live singlets across experimental timepoints, gated using a FMO (wild-type) control. **(C)** Live, lineage-negative, singlets plotted for EpCAM and CD49f to determine the distribution of luminal, basal, and stromal subpopulations across experimental timepoints. **(D)** Representative histograms of live singlets plotted based on GFP expression, split by luminal, basal, stromal, and lineage-positive fractions to highlight differences in GFP expression between populations and timepoints.
